# Chemical inhibition of bromodomain proteins in insect stage African trypanosomes perturbs silencing of the Variant Surface Glycoprotein repertoire and results in widespread changes in the transcriptome

**DOI:** 10.1101/2023.01.09.523188

**Authors:** Ethan C. Ashby, Jennifer L. Havens, Johanna Hardin, Danae Schulz

## Abstract

The eukaryotic protozoan parasite *Trypanosoma brucei, spp*. is transmitted by the tsetse fly to both humans and animals, where it causes a fatal disease called African trypanosomiasis. While the parasite lacks canonical DNA sequence specific transcription factors, it does possess histones, histone modifications, and proteins that write, erase, and read histone marks. Chemical inhibition of chromatin interacting bromodomain proteins has previously been shown to perturb bloodstream specific trypanosome processes, including silencing of the Variant Surface Glycoprotein genes (*VSG*s) and immune evasion. Transcriptomic changes that occur in bromodomain inhibited bloodstream parasites mirror many of the changes that occur as parasites developmentally progress from the bloodstream to the insect stage. We performed RNA-seq timecourses to determine the effects of chemical bromodomain inhibition in insect stage parasites using the compound I-BET151. We found that treatment with I-BET151 causes large changes in the transcriptome of insect stage parasites, and also perturbs silencing of *VSG* genes. The transcriptomes of bromodomain inhibited parasites share some features with early metacyclic stage parasites in the fly salivary gland, implicating bromodomain proteins as important for regulating transcript levels for developmentally relevant genes. However, the downregulation of surface procyclin protein that typically accompanies developmental progression is absent in bromodomain inhibited insect stage parasites. We conclude that chemical modulation of bromodomain proteins causes widespread transcriptomic changes in multiple trypanosome life cycle stages. Understanding the gene regulatory processes that facilitate transcriptome remodeling in this highly diverged eukaryote may shed light on how these mechanisms evolved.

## Intro

The African trypanosome *Trypanosoma brucei*, the causative agent of human and animal African trypanosomiasis, cycles between a mammalian host and a tsetse fly vector. Large differences in the two host environments necessitate extensive adaptation on the part of the parasite to ensure survival. This adaptation is facilitated by large changes in the transcriptome of parasites within each host ^1–6^. Parasites living in the bloodstream vary the proteins on their surface using a large repertoire of Variant Surface Glycoprotein (*VSG)* genes^7^ in order to evade the host immune system. Following transition to the fly after a bloodmeal, parasites remodel their surface proteins to express an invariant procyclin protein coded for by the *EP* and *GPEET* gene family^8^ and alter their metabolism to adapt to a glucose poor environment^9,10^. Within the fly, the parasites migrate from the midgut to the salivary gland, passing through a number of developmental stages before remodeling their surface proteins once more to prepare for entry into the mammalian bloodstream. The surface proteins expressed by these salivary gland metacyclic parasites are termed metacyclic VSGs (mVSGs)^11,12^. Because trypanosomes transition through a number of life cycle stages and diverged quite early compared to better studied model organisms, they serve as a model for gene regulatory mechanisms that facilitate transcriptome reprogramming and adaptation in early branching eukaryotes.

Trypanosomes have a number of unusual gene regulatory features, including polycistronic gene transcription and trans-splicing and polyadenylation of mRNAs^13,14^. Although trypanosomes lack canonical DNA sequence specific transcription factors, they do harbor histone modifications and the histone interacting enzymes required to write, erase, and read specific histone modifications^15,16^. Recent work has demonstrated that there are a large number of histone interacting proteins that form specific complexes in bloodstream forms, and that many of these complexes localize to sites of transcription initiation and termination^17^. Perturbation of histone interacting complexes in trypanosomes has been shown to affect immune evasion^18–23^, cell cycle^24^, and differentiation processes^23–25^.

We previously showed that small molecule and genetic inhibition of chromatin interacting bromodomain proteins in bloodstream parasites induces transcriptome reprogramming that shares similarities to transcriptomic changes that occur during the transition from the bloodstream to the procyclic form of the parasite^23^ in the insect midgut. Trypanosome bromodomain proteins bind to acetylated lysines^26,27^ and Bdf1-6^17,23,26^ have been shown to localize to transcription start sites. Another bromodomain protein (Bdf7) localizes to termination sites^17^. In addition, occupancy of the bromodomain protein Bdf3 is dynamic as parasites differentiate from bloodstream to procyclic forms, with occupancy peaking at 3h post differentiation^25^. The small molecule bromodomain inhibitor I-BET151 binds to TbBdf2 and TbBdf3, and may bind additional bromodomain proteins, though this has not been formally tested^23^. Treatment of bloodstream parasites with I-BET151 induces remodeling of the parasite surface wherein VSG proteins are downregulated and procyclin protein is expressed on the cell surface. In addition, large numbers of transcripts that are upregulated or downregulated during differentiation to the procyclic stage are similarly up or downregulated in I-BET151 treated bloodstream parasites^23^.

Accompanying the differentiation-like changes in I-BET151 treated parasites is a loss of monoallelic expression of *VSG* genes^23^. Immune evasion of the mammalian antibody response depends on the expression of only one VSG protein on the parasite surface at a time, and the rest of the large repertoire of *VSG* genes are silenced. I-BET151 treatment increases expression of *VSG* genes from silenced Bloodstream Expression Sites (BESs) and also increases the Expression Site Associated Genes (*ESAG*s) from these same sites. Metacyclic *VSG* genes and *VSG* genes at other sites in the genome are also derepressed in I-BET151 treated bloodstream parasites.

While the effect of small molecule bromodomain inhibition has been characterized in bloodstream parasites^23^, the effect of I-BET151 treatment in insect stage parasites is unknown. In order to further understand how bromodomain inhibition affects the transcriptome of insect stage parasites, we performed RNA-seq timecourses on I-BET151 treated midgut stage procyclic cells. Our analysis shows that large numbers of transcripts are altered in I-BET151 treated procyclic parasites and that drug treated parasites share some features with parasites in the early metacyclic salivary gland stages. We also show that silencing of *VSG* genes is perturbed in I-BET151 treated parasites, demonstrating that bromodomain proteins are important for gene silencing in multiple stages of the parasite life cycle.

## Results

### VSGs and ESAGs are among the most highly upregulated genes following bromodomain inhibition

In order to investigate the effects of bromodomain inhibition in insect stage parasites, we treated procyclic parasites with the previously validated bromodomain inhibitor I-BET151 ^23^. I-BET151 binds directly to Bdf2 and Bdf3, and may also bind other bromodomain proteins^23^. Procyclic parasites were treated with I-BET151 over the course of 2 weeks, and samples were harvested for RNA-seq at 10 timepoints (0h, 3h, 6h, 12h, 24h, 48h, 3d, 7d, 10d, 14d). We first performed a Principle Component Analysis (PCA) to confirm that biological replicates clustered together (Figure 1A). MA plots comparing DESeq normalized transcript levels at 12h of I-BET151 treatment with those of untreated parasites revealed a large number of genes with significantly altered transcript levels at this early timepoint (Figure 1C). We observed that transcript levels for *VSG* and *ESAG* genes were among the most highly upregulated in the dataset. This is consistent with previous work in bloodstream parasites showing that I-BET151 treatment and RNAi mediated silencing of individual bromodomains causes an increase in transcript levels of silent *VSG* genes^23^.

**Figure 1.**
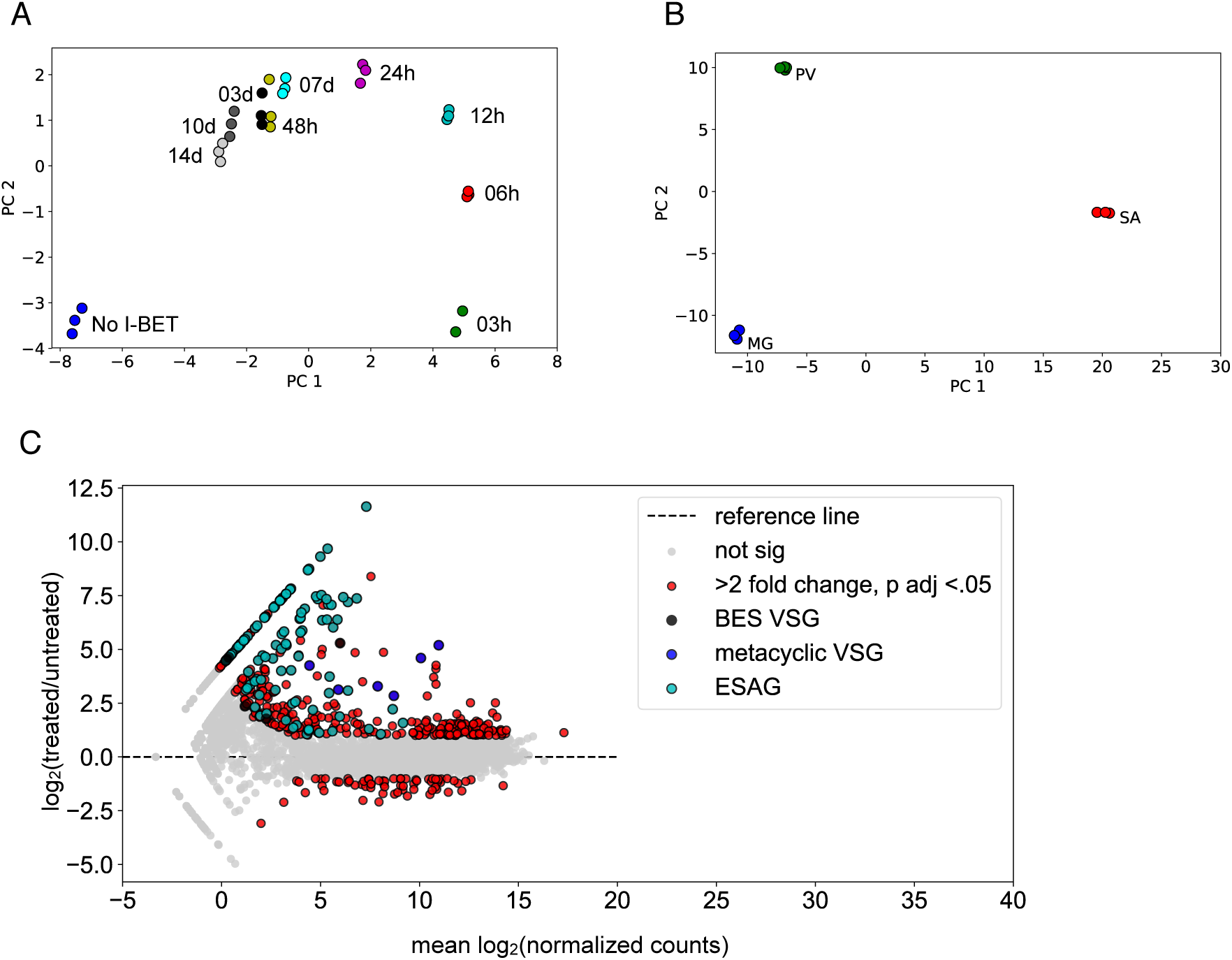
*VSG* and *ESAG* genes are the among the most highly upregulated genes following I-BET151 treatment in insect stage parasites. **A)** Principle component analysis of RNA-seq samples for I-BET151 treated parasites at indicated timepoints. **B)** Principle component analysis of RNA-seq samples from indicated tsetse fly organs using data from Savage et al.^28^. **C)** MA plot for parasites treated for 12 hours with I-BET151 compared with untreated parasites. *VSG* genes and *ESAG* genes are highlighted in black (BES *VSG*s), blue (metacyclic *VSG*s), and cyan (*ESAG* genes). Red dots indicate DESeq calculated differentially expressed genes with a Benjamini Hochberg adjusted p-value < 0.05 and a fold change > 2.

To further investigate the effect of I-BET151 treatment on *VSG* and *ESAG* transcript levels, we separated the *VSG* genes into subsets: VSGs within Bloodstream Expression Sites (BESs), metacyclic VSGs that contain specific promoters active in the salivary gland stage, and other VSGs that are located elsewhere in the genome. We plotted the median expression level for each of these sets of genes, and observed an increase in transcript levels for all of the VSG subsets and for *ESAG* genes at the 12h timepoint compared to untreated parasites (Figure 2A, 2B). This is the same phenotype as observed in bloodstream parasites following bromodomain inhibition^23^. We conclude from this data that bromodomain proteins are important for maintaining silencing of *VSG* transcripts in insect stage procyclic parasites in addition to their already established role in maintaining *VSG* silencing in bloodstream stage parasites.

**Figure 2.**
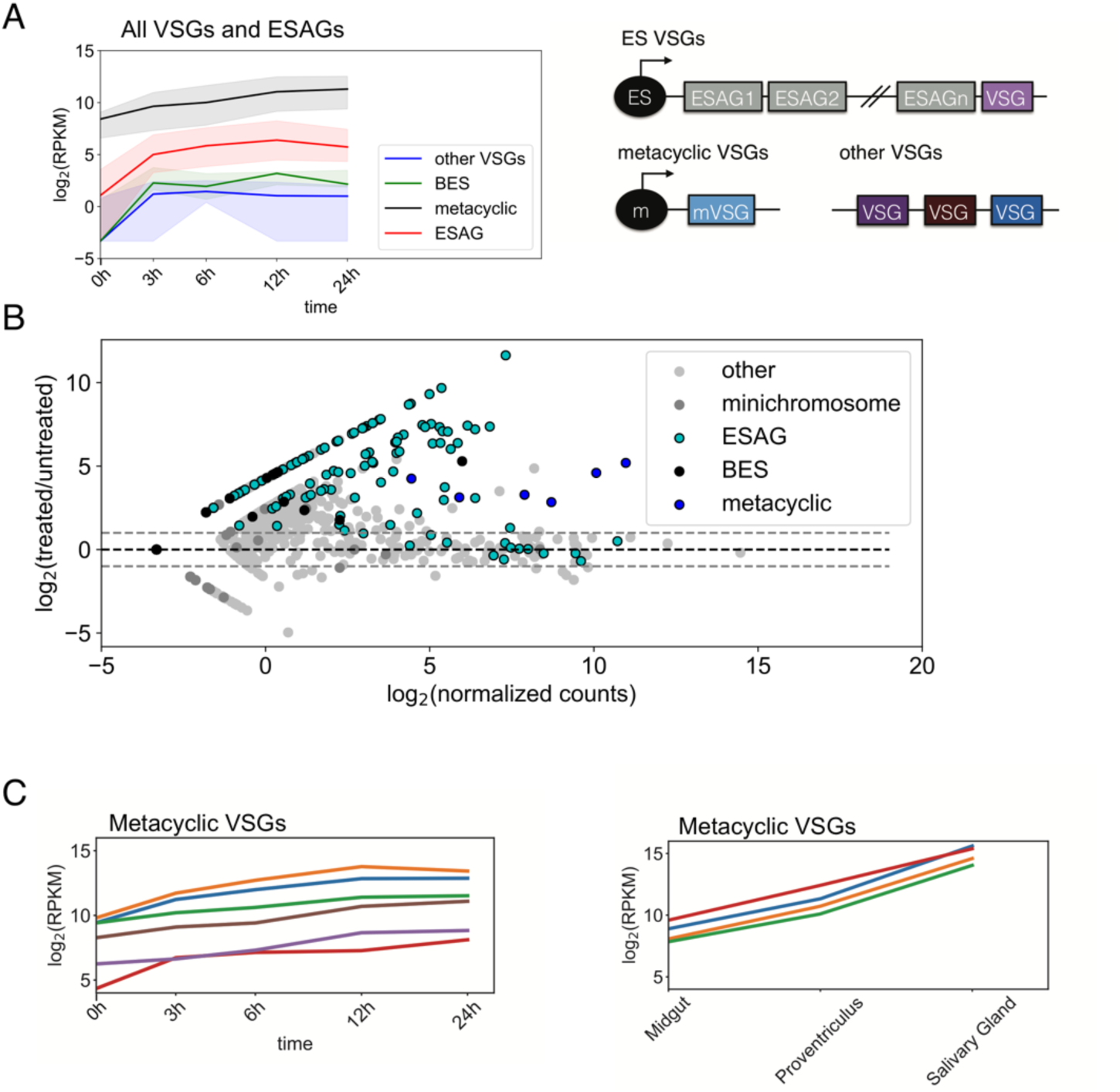
I-BET151 treatment in insect stage parasites increases transcript levels for *VSG* and *ESAG* genes that are silenced in wildtype parasites. **A)** Left, Median transcript levels for each indicated gene subset over 24h of I-BET151 treatment. Shading represents the inner quartile range at each timepoint. Right, Schematic of *VSG* gene locations in the genome. **B)** MA plot for *VSG* gene subsets and *ESAG* genes in parasites treated for 12 hours with I-BET151 compared with untreated parasites. Dashed lines demarcate 2-fold differences in treated parasites compared to untreated parasites. **C)** Left, Transcript levels for 6 individual *VSG* genes with metacyclic promoters over 24h of I-BET151 treatment. Right, Transcript levels for 4 *VSG* genes in samples taken from indicated organs in the tsetse fly using data from Savage et al^28^.

### Metacyclic *VSG* transcript levels are increased following I-BET151 inhibition

Bromodomain inhibition in bloodstream parasites results in changes in the transcriptome that resemble many of the changes in transcript levels that occur as bloodstream parasites transition to the procyclic stage^23^. We wanted to test the model that bromodomain inhibition might initiate a default program that results in transcript changes consistent with developmental progression. This model predicts that inhibition of bromodomain proteins in gut stage procyclic parasites might cause transcriptome changes similar to those that occur as gut stage parasites migrate to the proventriculus and from there to the salivary gland. To investigate this question, we took advantage of an existing RNA-seq dataset generated from parasites living in the fly gut, proventriculus, and salivary gland^28^, which we will henceforth refer to as the insect stage development dataset. We obtained fastq files from the insect stage development dataset and analyzed them using our RNA-seq pipeline to ensure the dataset was comparable to the bromodomain inhibition dataset that we generated. A PCA analysis demonstrated that the replicates from the insect stage development dataset clustered together following analysis with our pipeline (Figure 1B). One of the most prominent changes in gene expression following transition to the salivary gland is the upregulation of *VSG* genes with metacyclic promoters that facilitates transition to a bloodstream environment^29^. We plotted the expression of *VSG* genes with metacyclic promoters following I-BET151 treatment, and compared it to the expression of metacyclic *VSG* genes in the insect stage development dataset (Figure 2C). As expected, we observed that the expression of metacyclic *VSG* genes increased sharply as parasites transitioned to the salivary gland. We also observed an increase in transcript levels of metacyclic *VSG* genes following I-BET151 treatment (note that the metacyclic VSG genes are named differently in the two different strains used) (Figure 2C). This data indicates that I-BET151 treatment and developmental progression have similar effects on transcript levels for the metacyclic *VSG* gene subset. However, since most *VSG* subsets show increased transcript levels following I-BET151 treatment, we also investigated the behavior of other sets of genes known to be associated with developmental progression.

### Some gene sets associated with developmental progression show changes in transcript levels following I-BET151 treatment

We next investigated whether other gene sets identified in the insect stage development dataset were also altered following treatment with I-BET151. Savage et al. identified a number of gene sets with altered transcript levels as parasites progressed from the midgut to the salivary gland. These included a set of adenylate cyclases, transporters, RNA binding proteins, and surface proteins^28^. We separated each gene set into those that increased during developmental progression and those that decreased. We performed Gene Set Enrichment Analysis (GSEA) to see if any of these gene sets were overrepresented in the set of differentially expressed genes following treatment with I-BET151. Genes with transcript levels that change early in the timecourse are likely a more direct result of bromodomain inhibition than changes that occur later, so we focused on genes that were differentially expressed by 12 hours following bromodomain inhibition. Of the gene sets tested this way, the set of adenylate cyclases and transporters that were downregulated during developmental progression also showed downregulation following bromodomain inhibition (Figure 3) and showed significant enrichment in the GSEA analysis (Supplemental Table 1).

**Figure 3.**
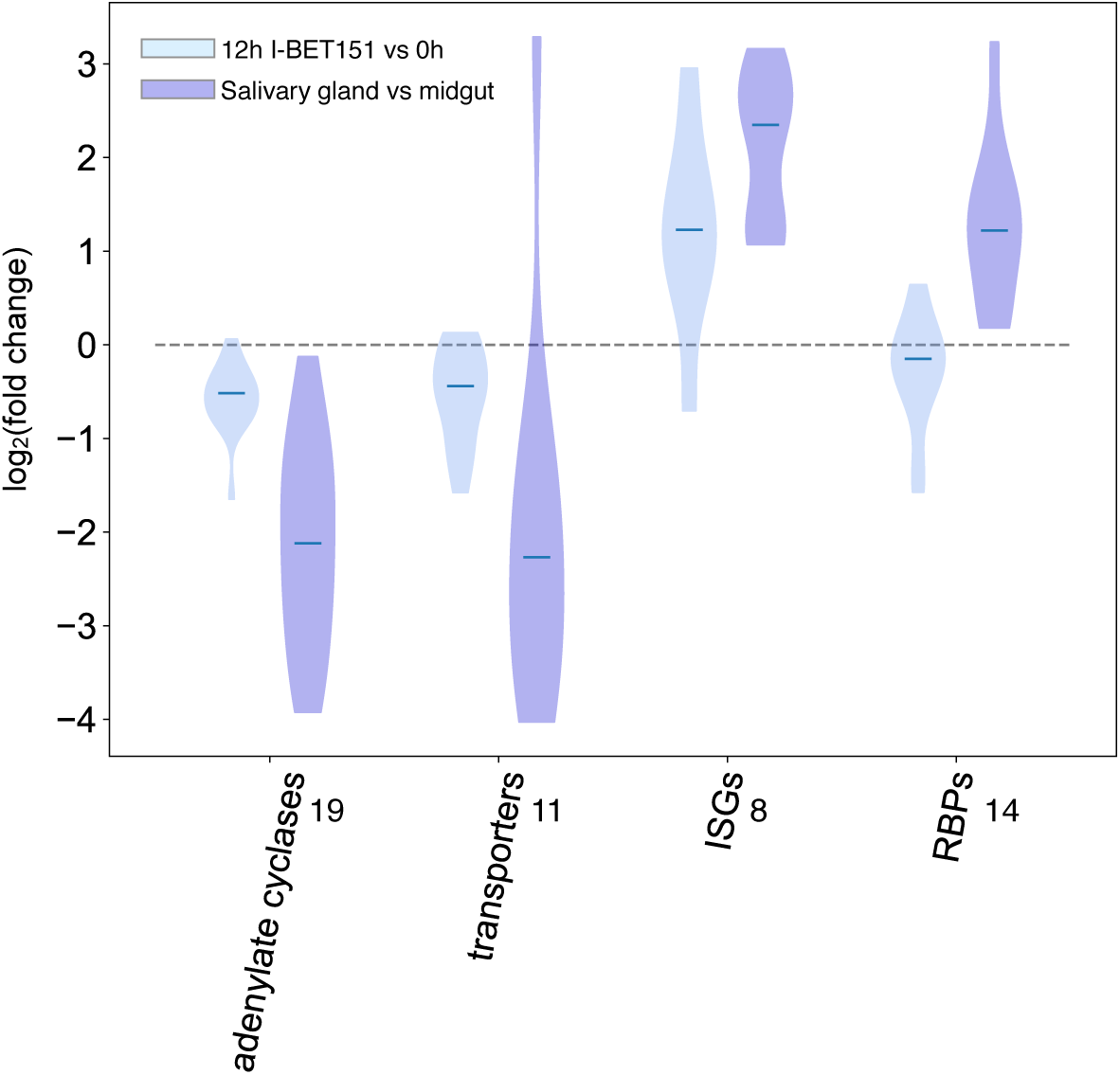
Transcript levels for some gene sets associated with insect stage developmental progression are also altered following I-BET151 treatment. Violin plot for transcript levels of adenylate cyclases and transporters that are downregulated during insect stage developmental progression and transcript levels of *ISG*s and *RBP*s that are upregulated during developmental progression. Data are plotted as the log_2_(I-BET151 (12h)/untreated (0h)) and log_2_(salivary gland/midgut) using data on parasites harvested from the midgut and salivary gland from Savage et al^28^. All gene sets shown were statistically significant using GSEA (Supplemental Table 1). Number to the right of the x-axis label indicates number of genes in the gene set analyzed.

In addition, many of the invariant surface glycoproteins (ISGs) that were identified as upregulated in the insect stage development dataset were also upregulated following I-BET151 treatment (Figure 3). Although upregulated RBPs from the Savage et al. dataset were significantly enriched by GSEA in I-BET151 treated parasites, some members of this set increased while others decreased, and thus no clear cut pattern for expression emerged for this group (Figure 3). Thus, bromodomain inhibition in procyclic parasites results in some transcript level changes that are similar to those that occur during insect stage developmental progression. However, we note that patterns of gene expression that are considered hallmarks of the transition from parasites living in the gut to those in the salivary glands do not show significantly altered levels in I-BET151 treated procyclic parasites. Specifically, the *EP* genes that are downregulated and the *BARP* genes that are upregulated during the transition to the epimastigote stage do not show significantly altered transcript levels following I-BET151 treatment. We verified that procyclin expression was not altered in procyclic parasites following I-BET151 treatment using a previously validated *EP1/GFP* reporter parasite line^30^ (Supplemental Figure 1). We observed no difference in *EP1/GFP* expression in I-BET151 treated parasites (Supplemental Figure 2).

### Parasites treated with I-BET151 share transcriptomic features with early metacyclic parasites

We next compared I-BET151 induced transcriptomic differences with those described for a single-cell sequencing dataset cataloguing developmental progression in insect stage parasites^31^. The advantage of this dataset is that the metacyclic stages are separated into more subsets than in the Savage et al. study. Vigneron et al.^31^ separated parasites into 3 broad categories: epimastigote, meta 1, and meta 2, with meta 1 representing early metacyclic parasites and meta 2 representing later stage metacyclics. Each broad category was selected for a set of biomarker genes that represented highly differentially expressed genes. We performed GSEA for these 3 developmental group biomarker genes on our I-BET151 dataset. While all gene groups were significantly enriched in our I-BET151 differentially expressed dataset, the early metacyclic Meta 1 biomarker gene set was specifically upregulated (Figure 4A, Supplemental Figure 3). Vigneron et al. also catalogued the expression of a set of signature genes during developmental progression. These include a set of RNA Binding Proteins (RBPs), Fam50 and Lipid Phosphate Phosphatase (LPP) genes, Fam10 genes, and calpains. With the exception of the calpains, all these gene sets were significantly enriched in our I-BET151 treated parasites by GSEA (Supplemental Table 1). Although the calpain genes did not meet our FDR threshold for GSEA analysis, visual inspection of this gene set showed an increase in transcript levels for all 3 genes, which peak in expression at the early Meta 1 stage in developing parasites. Vigneron et al. found that a set of RBP genes peak at the meta 1 stage, while the expression of the Fam50 and LPP gene set decrease at the meta 1 stage. I-BET151 treated parasites also showed increases in these same RBP genes, while the Fam50 and LPP gene sets briefly increased and then decreased by 12h (Figure 4A and 4B). While Vigneron et al. observed an increase in transcript levels for Fam10 genes in late meta1 and early meta 2 stages, we did not see an upregulation of that gene set in I-BET151 treated parasites.

**Figure 4.**
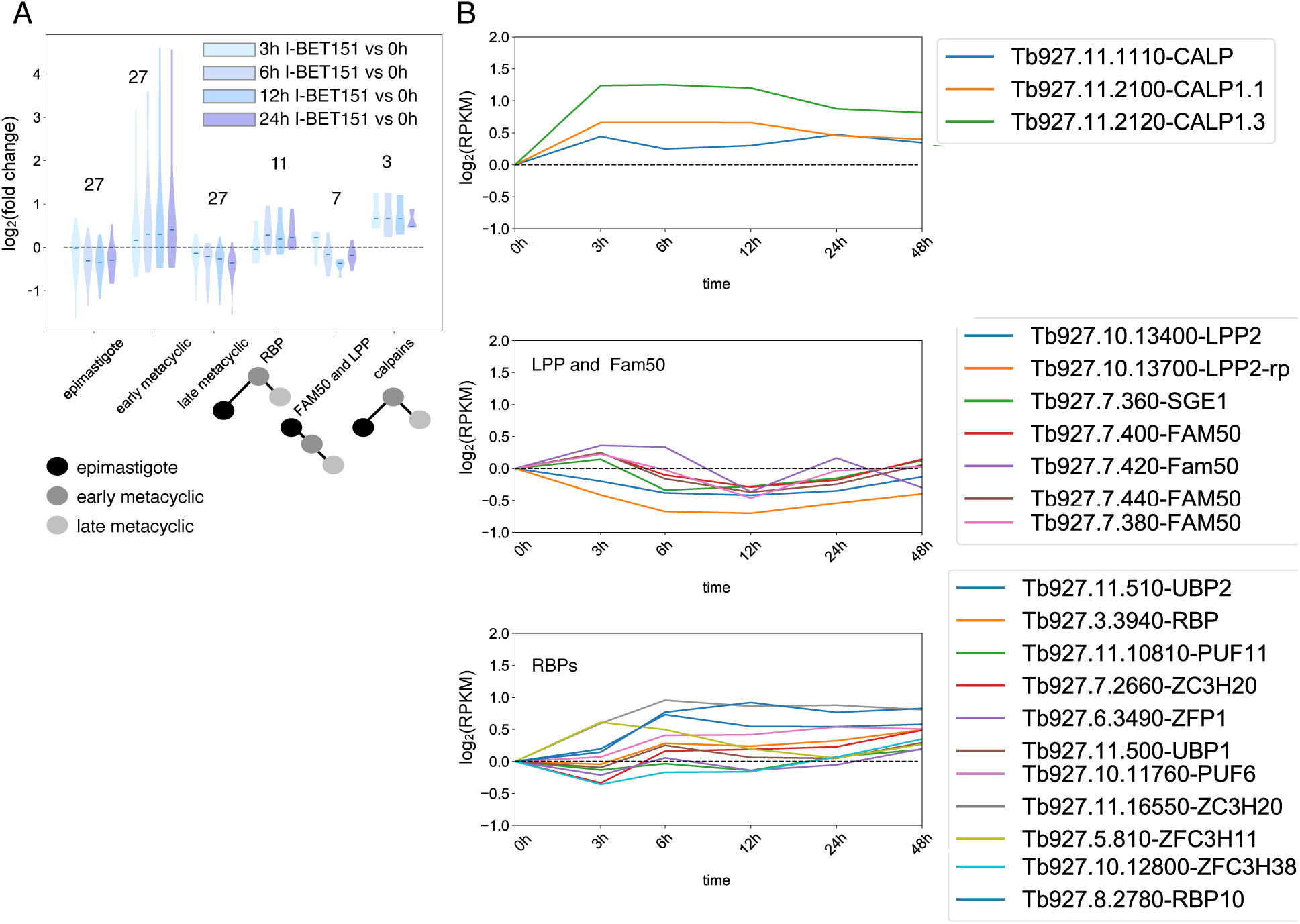
I-BET151 treated parasites share some transcriptomic features of early metacyclic parasites. **A)** Violin plot showing transcript levels of indicated gene sets defined by Vigneron et al^31^ in parasites over 24 hours of I-BET151 treatment. Number over gene set indicates number of genes in set. Data are plotted as the log_2_(I-BET151 treated/untreated (0h)) using normalized counts. Blue shading indicates time of I-BET151 treatment. Schematic under label represents transcript levels for each group during development as measured by Vigneron et al.^31^, where a black node indicates transcript level at the epimastigote stage, a dark gray node represents transcript level in early metacylic parasites, and light gray node indicates transcript level in late metacyclic parasites. **B)** Normalized transcript levels for the indicated genes following I-BET151 treatment. Each gene set was identified by Vigneron et al.^31^ as having altered expression specifically in early metacyclic stage parasites.

Overall, we conclude that a model wherein bromodomain inhibition leads to a default program of developmental progression is too simplistic to describe the observed data. Our analysis shows that I-BET151 treated parasites share transcriptomic features with early metacyclic parasites, including upregulation of the Meta 1 biomarker gene set (Figure 4A) and changes in gene expression for adenylate cyclases, transporters, RBPs, calpains, LPP, and Fam50 genes. However, some hallmarks of developmental progression are not observed, including upregulation of *BARP*, downregulation of the *EP* genes, and upregulation of the Fam10 gene set, which has been shown to occur in the transition from early to late metacyclic stages^31^.

### Clustering analysis reveals that differentially expressed genes in I-BET151 treated parasites can be grouped into 6 distinct expression patterns

We next analyzed the I-BET151 dataset to identify groups of genes whose transcript levels change with similar timing and magnitude. As we were particularly interested in early effects following bromodomain inhibition, we identified all differentially expressed genes across 5 timepoints in the first 24 hours following I-BET151 treatment (0h, 3h, 6h, 12h, 24h) using a likelihood ratio test (LRT). This analysis revealed that a large number of 6548 genes were differentially expressed in the first 24h after bromodomain inhibition with a Benjamini-Hochberg p adjusted value < .01. We then used the Mfuzz program^32^ to group each of these genes into one of six clusters of genes (Figure 5). We identified three clusters of upregulated genes (Clusters 1,3, and 6) and three clusters of downregulated genes (Clusters 2,4, and 5) (Figure 5). Clusters 1 and 2 have a sharp change in expression at 3h that rapidly reverses toward the starting transcript levels. Clusters 5 and 6 have sharp changes in expression at 3h and this change persists throughout the first 24h of I-BET151 treatment. Finally, Clusters 3 and 4 have more gradual changes in expression levels that also persist for the first 24h. Of all the upregulated *VSG* genes that were observed, the majority fell within Cluster 3, which shows a more gradual increase in gene expression compared to Clusters 1 and 6. We conclude that bromodomain inhibition in insect stage parasites results in large changes in gene expression, many of which occur quite early within 6 hours of treatment.

**Figure 5.**
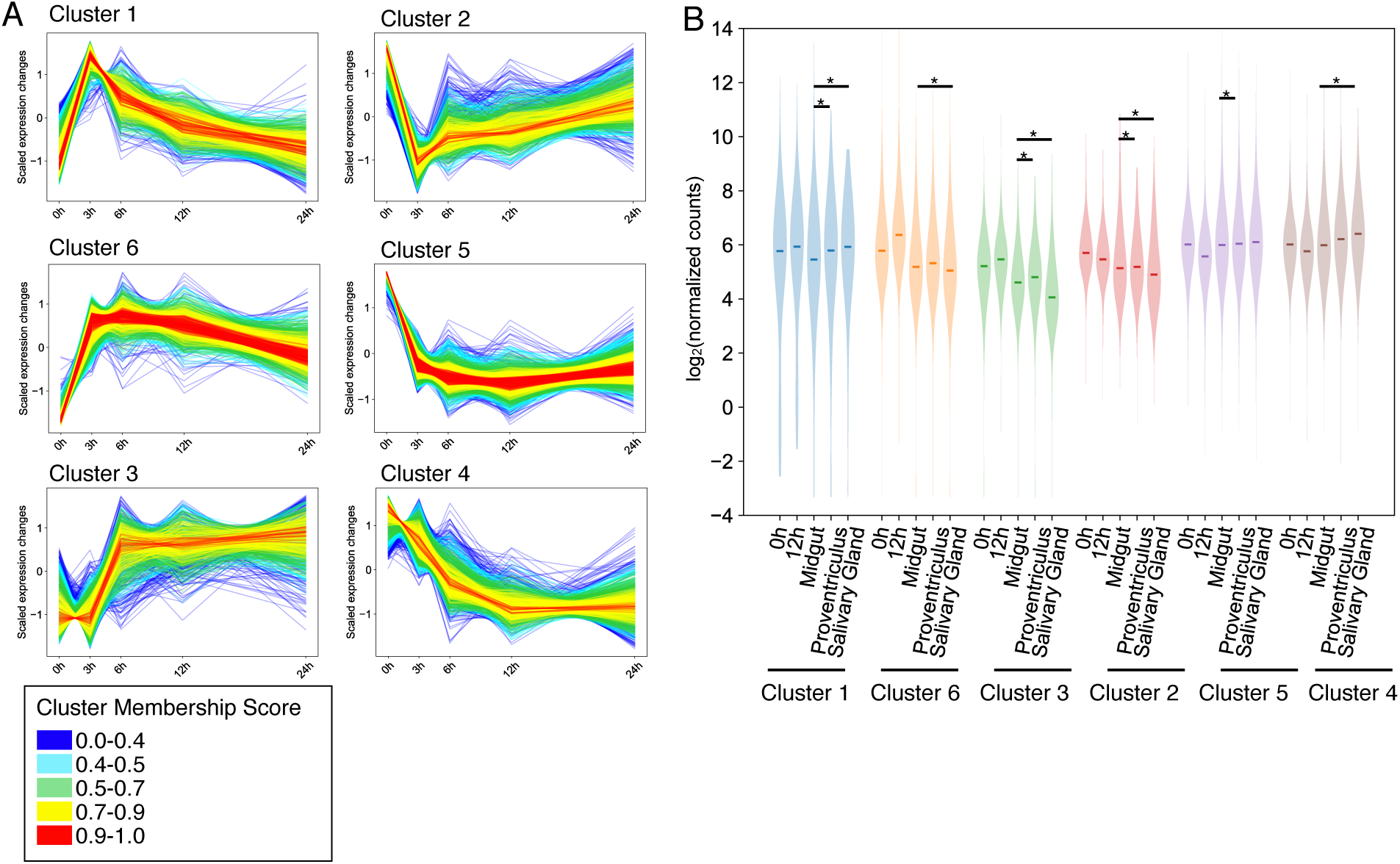
Differentially expressed genes in I-BET151 treated parasites show distinct patterns of expression. **A)** DESeq normalized transcript levels of differentially expressed genes following I-BET151 treatment and clustered based on expression pattern and timing. Transcript levels are scaled such that the mean is 0 and the standard deviation is 1. **B)** Violin plot comparing changes in transcript levels for each cluster in **A)** following 12h of I-BET151 treatment or during insect stage developmental progression using data for midgut, proventriculus, and salivary gland parasites taken from Savage et al^28^. Stars indicate cluster identified as enriched by GSEA with an FDR < 0.1 in the insect stage developmental progression dataset.

We next looked at whether the clusters of genes we identified following I-BET151 treatment were enriched in the differentially expressed genes from the insect stage development dataset (Savage et al.). To do this we first filtered each cluster for a membership correlation value of 0.7, which resulted in 2,549 genes spread out among the 6 clusters (Supplemental Table 6). We then interrogated the insect stage development dataset to see if these high membership clustered gene sets were overrepresented in parasites developing in the proventriculus or the salivary gland. All the high membership clusters were significantly overrepresented by GSEA in parasites harvested from either the proventriculus or the salivary gland compared to midgut parasites (Supplemental Table 1). However, only 4 clusters showed changes in I-BET151 treated parasites consistent with the direction of change during developmental progression. Specifically, Clusters 1, 3 and 6 show median expression values that increase with I-BET151 treatment; an increase in median expression values for these clusters is also observed as parasites progress from the midgut to the proventriculus. Cluster 1 shows median expression values that increase with I-BET151 treatment; an increase in median expression value is also observed for this cluster as parasites progress from the midgut to the salivary gland. Finally, a decrease in median expression value is observed for genes in Cluster 2 following I-BET151 treatment, and a decrease in expression value is also seen for this cluster as parasites progress from the midgut to the salivary gland.

We performed a GO analysis on each of the high membership clusters we identified in the I-BET151 treatment dataset. We filtered the GO terms using a Benjamini-Hochberg adjusted p-value of less than .05 and an enrichment score greater than 3 (Supplemental Table 4). While this analysis was not revealing for many of the clusters, Cluster 3 was particularly enriched for GO terms having to do with cell movement, particularly with microtubule based processes (Figure 6 and Supplemental Figure 4). This is consistent with a Meta 1 developmental phenotype as defined by Vigneron et al., who found that genes associated with cell movement were particularly upregulated at this developmental stage^31^. Cluster 3 is more slowly upregulated following bromodomain inhibition and the upregulation persists for the first 24h of treatment (Figure 6A). We plotted the expression levels for genes within each GO enriched group for Cluster 3 following bromodomain inhibition and during insect stage developmental progression (Figure 6B). These results demonstrate that the median expression level for genes associated with movement in Cluster 3 increases following I-BET151 treatment and as parasites transition from the midgut to the proventriculus. We conclude that bromodomain inhibition causes changes in transcript levels for genes associated with microtubule related processes important for cell movement, which are also processes involved in developmental progression.

**Figure 6.**
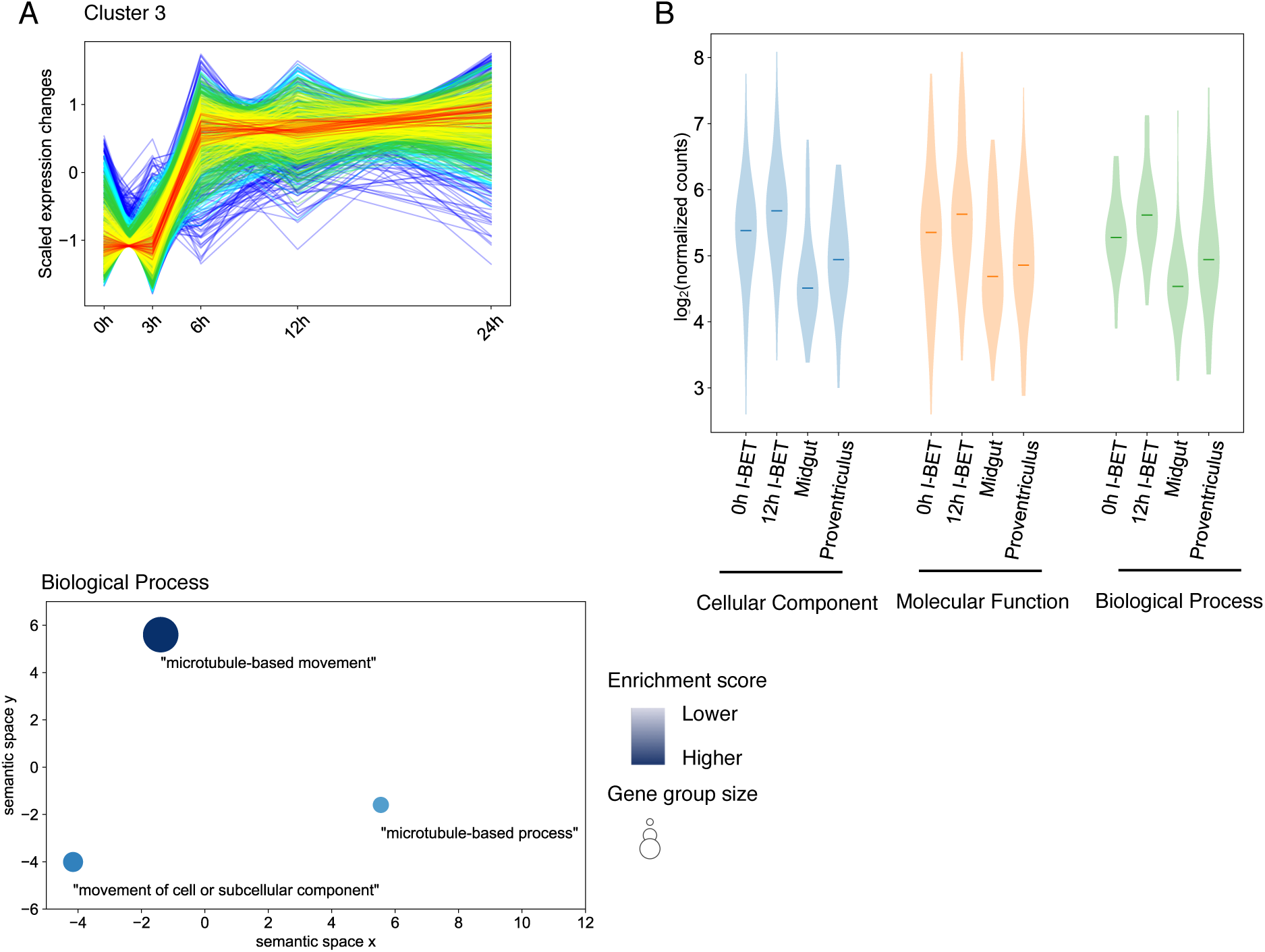
Genes involved in movement and microtubule based processes are upregulated in I-BET151 parasites and in parasites transitioning from the midgut to the proventriculus. **A)** Top, Same as in **Figure 6A.** Bottom, REVIGO^80^ plot of GO enrichment analysis performed on Cluster 3. **B)** Violin plot for genes identified as enriched by GO analysis of Cluster 3. Transcript levels for I-BET151 treated parasites are compared with transcript levels in midgut and proventriculus stage parasites using data from Savage et al^28^.

### Gene ontology analysis reveals that I-BET151 treatment causes changes in transcript levels for genes associated with a wide array of cellular processes

To better capture the full array of transcriptome changes that occur following I-BET151 treatment in insect stage parasites, we performed GSEA on sets of genes grouped by Gene Ontology (GO). In this analysis, we once again focused on genes that were differentially expressed between the 0h timepoint and 12h of I-BET151 treatment. This analysis revealed that a large number of gene sets were affected. Biological processes associated with antigenic variation and evasion of the host immune response, adenylate cyclase activity, translational initiation, rRNA processing, intracellular signal transduction, translation initiation factor activity, cAMP biosynthesis, nucleobase containing compound metabolic processes, and cyclic nucleotide biosynthetic processes were all significantly enriched in the differentially expressed gene set following 12h of bromodomain inhibition (Supplemental Table 1 and Figure 7). The large number of gene sets affected by I-BET151 treatment indicate that bromodomain inhibition has effects on a diverse set of biological processes, and that the transcriptomic differences observed following treatment likely reflect direct and indirect effects of the drug, even at early timepoints.

**Figure 7.**
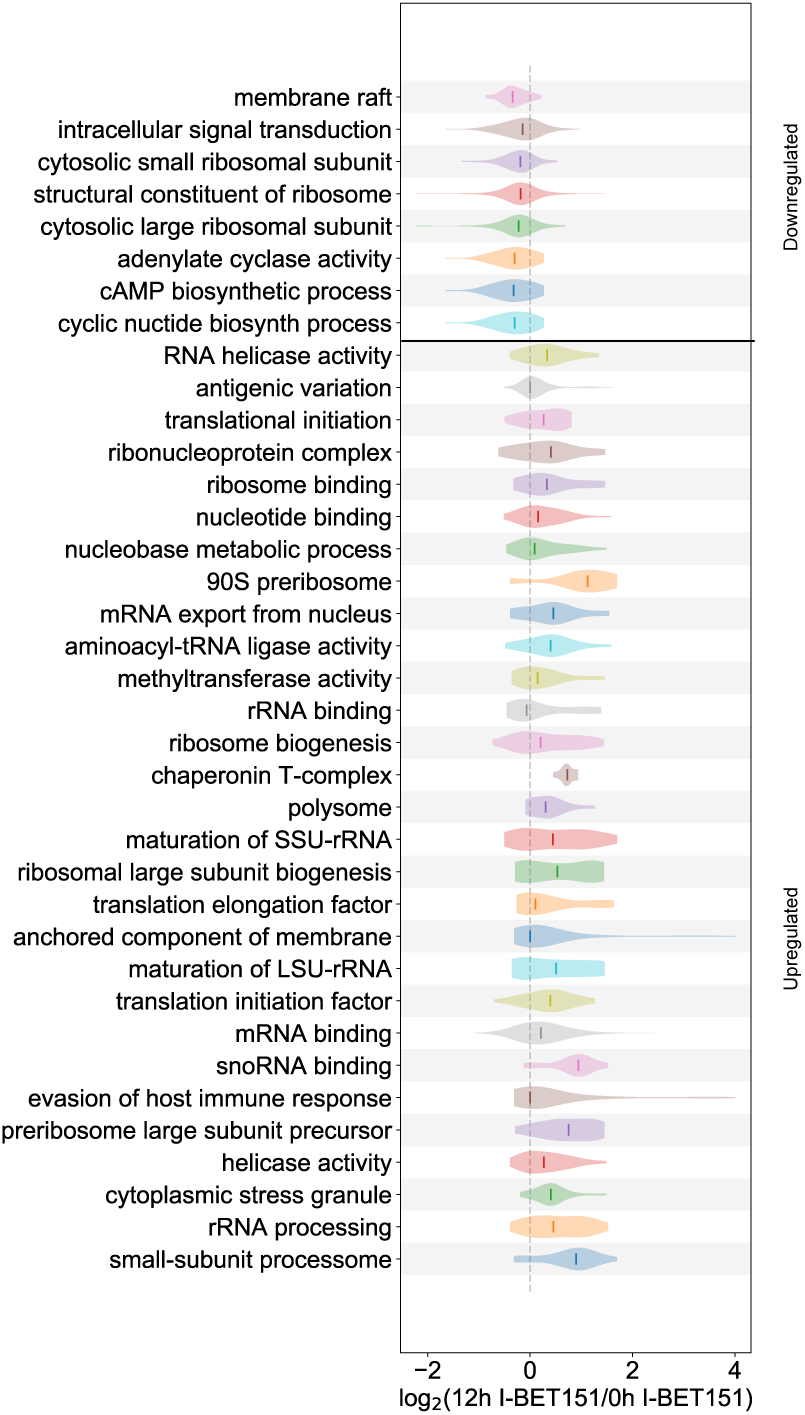
Diverse biological processes affected by I-BET151 treatment in insect stage parasites. Violin plot showing transcript levels of parasites treated for 12h with I-BET151 compared with untreated parasites (0h) parasites expressed as log_2_(12h I-BET151/0h IBET151) using normalized counts for GO sets with an FDR of < 0.1 by GSEA.

### I-BET151 driven transcriptome changes vary for different life cycle stages

To try to understand whether the effects of bromodomain inhibition are context dependent, we compared the most highly upregulated and downregulated genes following bromodomain inhibition in bloodstream form parasites and procyclic form parasites. To do this, we took advantage of a previously published RNA-seq timecourse on bloodstream form parasites treated with I-BET151^23^. We used DESeq2 to ascertain which differentially expressed genes had altered transcript levels of >2-fold up or down at any time point compared to untreated parasites. We separated transcripts into those that were increased or decreased following bromodomain inhibition and generated an upset plot to ascertain which genes were in common between the bloodstream dataset and the procyclic dataset (Figure 8). This analysis revealed that a large number of transcript levels are upregulated or downregulated only in one of the life cycle stages analyzed. Of the 746 transcripts that showed large changes following bromodomain inhibition, only 75 were affected by I-BET151 treatment in both procyclic and bloodstream forms. Of these, the largest group were 44 genes whose transcript levels are downregulated in bloodstream forms and upregulated in procyclic forms following I-BET151 treatment (Supplemental Table 5). This group includes 6 *ISG*s, which are downregulated during differentiation from bloodstream to procyclic forms^5^ and upregulated in salivary gland parasites^28^.

**Figure 8.**
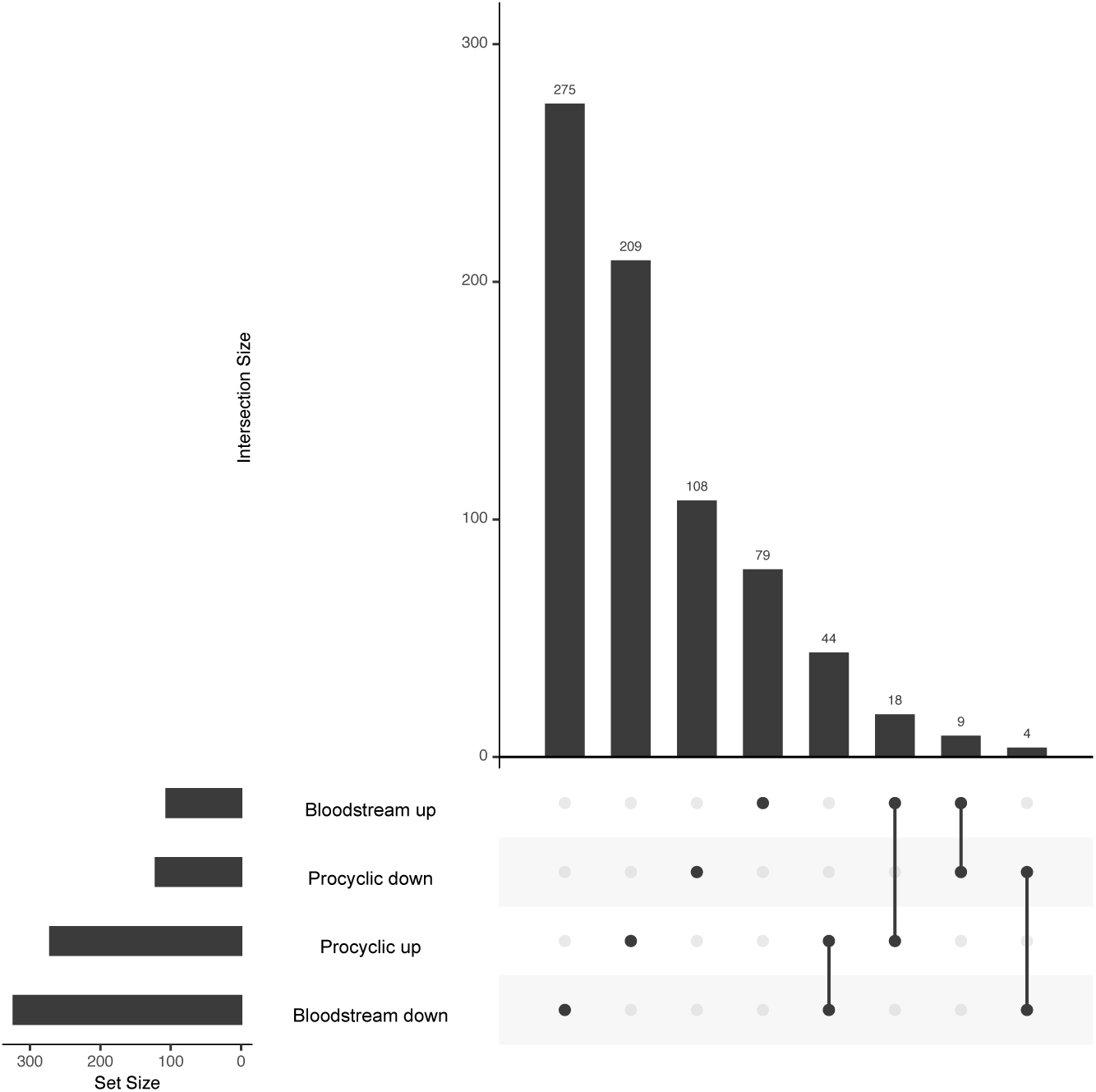
Many transcriptomic changes initiated by I-BET151 treatment are life cycle stage specific. Upset plot showing differentially expressed genes identified by DESeq with a Benjamini-Hochberg adjusted p-value < 0.05 and fold change > 2 up or down in bloodstream and procyclic stage parasites. Data from bloodstream parasites is generated from Schulz et al.^23^.

The next largest group of 18 genes with large changes in both bloodstream and procyclic bromodomain inhibited parasites were upregulated in both forms. This group included 4 *PAG*s, 1 Protein Associated with Differentiation (*PAD3*), and a number of other genes that are highly upregulated in parasites transitioning from the bloodstream form to the procyclic form (Tb927.8.520, Tb927.8.480, Tb927.3.590, Tb927.10.9550, Tb927.11.4700, Tb927.2.3460, Tb927.8.1270, Tb927.3.2750, Tb927.10.2310)^5^. In this group, an adenosine transporter (Tb927.3.590), a prostaglandin F synthase (Tb927.11.4700), and a cysteine peptidase family gene (Tb927.2.3460) were highly upregulated in parasites moving from the midgut to the salivary gland. 9 genes were upregulated in bromodomain inhibited bloodstream forms and downregulated in procyclic forms, while 4 genes were downregulated in bloodstream forms and upregulated in procyclic forms. We conclude that the effect of I-BET151 on the transcriptome is highly context dependent on life cycle stage. With the exception of the *VSG*s and *ESAGs*, the sets of genes that are upregulated or downregulated following I-BET151 treatment in procyclic vs bloodstream forms do not contain large overlaps (Figure 8).

## Discussion

Bromodomain inhibitors have garnered quite a bit of attention in recent years because of their potential as anti-cancer therapeutics. BET inhibitors have been shown to inhibit tumor growth in a number of models, including Acute Myeloid Leukemia (AML) ^33–37^, prostate cancer^38^, neuroblastoma^39^, and breast cancer^40^. In parasites, bromodomain inhibitors have been shown to bind to bromodomain proteins in *T. brucei, T. cruzi, P. falciparum*, and *L. dovani* ^23,27,41–44^. Inhibition of bromodomain proteins in parasites has been shown to affect differentiation processes in multiple parasite systems ^23,45,46^, and therapeutic strategies targeting chromatin interacting proteins have also been proposed and/or demonstrated for trypanosomiasis ^23,30^, Chagas disease ^47–49^, schistosomiasis^50,51^, toxoplasmosis ^52,53^, leishmaniasis^54^, and malaria ^55,56^.

While the biological effects of the bromodomain inhibitor I-BET151 have been documented for bloodstream stage *T. brucei* parasites^23^, nothing is known about the effect of bromodomain inhibition by this molecule at other parasite life cycle stages. The work presented here indicates that perturbations in *VSG* silencing following treatment with I-BET151 occurs in both bloodstream and insect stage parasites (Figure 2), whereas the effects on Pol II-driven genes are more context dependent and depend on the life cycle stage (Figure 8). I-BET151 inhibition in bloodstream parasites causes remodeling of cell surface proteins, wherein I-BET151 treated parasites show a decrease in VSG protein on the surface and a concomitant increase in procyclin protein^23^. While metacyclic *VSG* genes were upregulated in I-BET151 treated insect stage parasites, we detected no differences in the expression of procyclin (Supplemental Figure 2), indicating that extensive remodeling of the cell surface is unlikely to occur following bromodomain inhibition. TbSAP1 was recently identified as an important repressor of metacyclic VSG expression^22^. We observed reasonably high and unvarying transcript levels of TbSAP1 following bromodomain inhibition, and it’s possible that the presence of this protein prevented sufficient expression of metacyclic *VSG* genes to support surface remodeling. Alternatively, additional checkpoints might be required to remodel the cell surface that were not induced by treatment with I-BET151. In bloodstream parasites, loss of *VSG* silencing has been tied to life cycle progression^57,58^. In parasites undergoing metacyclogenesis, all metacyclic *VSG* genes are upregulated prior to commitment to one particular metacyclic VSG^59^. However, because the VSGs have not been as extensively catalogued in the parasite strain used for the insect developmental progression dataset, it is unclear whether loss of VSG silencing for non-metacyclic *VSG* genes is observed during insect stage developmental progression.

I-BET151-treated procyclic parasites share some features with insect stage differentiating parasites. I-BET151-treated parasites downregulate a set of adenylate cyclases that are also downregulated in salivary gland parasites (Figure 3). These include flagellar receptor adenylate cyclases *ACP3-5* (*Tb927*.*10*.*13040, Tb927*.*10*.*13740, Tb927*.*7*.*7470*) and a large number of *GRESAGs*. ACP3-5 are procyclic stage specific flagellar proteins^60^, and other members of this family (ACP1, 2 and 6) have been shown to be important for social motility^61^. ACP2 (Tb927.10.16190) and ACP6 (Tb927.9.15660) were also differentially expressed in our I-BET151 treated parasites. Salivary gland parasites also upregulate invariant surface glycoproteins (*ISG*s)^62,63^ in preparation for transition to the bloodstream stage, and we found this set of genes was also upregulated in I-BET151 treated parasites. The expression of *ISG* genes has recently been shown to be important for surviving the complement defense system in the mammalian bloodstream^64^. Several glucose transporters important for the glycolytic pathway are also downregulated as parasites differentiate within the insect, and we found that the THT1E (Tb927.10.8450 THT1E) and THT1-glucose transporters (Tb927.10.8440, Tb927.10.8470) were downregulated in I-BET151 treated parasites. Other transporters, such as nucleobase transporter Tb927.11.3620, cation transporter Tb927.11.8990, and pteridine transporter Tb927.10.9080 are downregulated in differentiating and I-BET151 treated parasites. This may reflect the differing metabolic requirements of parasites at different stages within the fly compartment.

The elegant single cell studies of Vigneron et al. delineated key biomarkers for parasite development within the fly. Of these classified biomarkers, we found that the early metacyclic Meta1 biomarkers were most upregulated in I-BET151 treated parasites (Figure 4). Several families of genes defined by Vigneron et al.^31^ showed similar expression patterns at the early metacyclic stage and in I-BET151 treated parasites, including a set of 11 RBPs, the Fam50 genes^65^ (Tb927.7.360, Tb927.7.400, Tb927.7.420, Tb927.7.440, Tb927.7.380), the LPP genes Tb927.10.13400 and Tb927.10.13700, and the calpain genes (Tb927.11.1110, Tb927.1.2100, Tb927.1.2120). Interestingly SGE1, a member of the Fam50 gene family, has been shown to be upregulated in epimastigotes but downregulated thereafter^31^. We find that SGE1 and other Fam50 family members in I-BET151 treated parasites are transiently upregulated at early timepoints, and then downregulated thereafter (Figure 4). However, the set of Fam10 SGM proteins that are most upregulated at late stage Meta 1 and Meta 2 stages of development are not upregulated in I-BET151 treated parasites. This supports the idea that I-BET151 treatment might initiate some features of the insect stage developmental program, but that other features are under control of other gene regulatory forces. SGM 1.7 has been definitively localized to the cell surface, and it’s possible that surface protein remodeling requires additional checkpoints that are not initiated with I-BET151 treatment, as discussed above. Interestingly, the set of Cluster 3 genes that peaks at 6h in the I-BET151 dataset (Figure 6) is enriched in microtubule based processes, and this is also a GO term set that is enriched in the early Meta 1 parasites^31^.

Independent of the effect of I-BET151 treatment on developmental genes, it’s clear from the cluster analysis that I-BET151 treatment results in large global changes in gene expression, and that these changes include both upregulation and downregulation (Figure 5). In addition, the timing and pattern of the expression changes varies from early changes at 3h that rebound quickly (Clusters 1 and 2) to slower changes that are more sustained (Clusters 3 and 4). Given the observation that large numbers of genes show similar expression patterns, we favor a model wherein I-BET151 treatment influences the expression of one or a few RNA binding proteins that themselves regulate large numbers of genes through interaction with their 3’ UTRs^66,67^. RNA binding proteins have been shown to be critical for postranscriptional regulation in trypanosomes (reviewed in^68^) and have also been implicated in differentiation processes in both bloodstream^69–72^ and insect stage forms^73–76^. Indeed we find that 8 of the 11 RBPs reported by Vigneron to be differentially expressed during insect stage development were differentially expressed in our dataset, including RBP10 (Tb927.8.2780), which is upregulated 2 fold after I-BET151 treatment and also upregulated at early metacyclic stages^31^. One reason why some features of insect stage differentiation, such as surface protein expression, may not be recapitulated in I-BET151 treated parasites is that one or more RBPs important for this process are not activated following drug treatment. This would be consistent with a model wherein different RBPs may be responsible for different aspects of the differentiation program, and thus it may be possible to induce some parts of the program and not others.

While both TbBdf2 and TbBdf3 have been shown to bind I-BET151 *in vitro*^23^, it remains unknown whether this molecule can bind any of the other five predicted bromodomain proteins in *T. brucei*. Elegant work by Staneva et al has shown that while 6 of the 7 predicted trypanosome bromodomain proteins localize to transcription start sites, these proteins do not exist all together in a single complex^17^. Thus, trypanosome bromodomain proteins likely have redundant but not completely overlapping functions, and it would be interesting to ascertain if other trypanosome bromodomain proteins bind to I-BET151. While many of the effects of I-BET151 in bloodstream parasites can be recapitulated by individual knockdown of TbBdf3, it remains possible that I-BET151 could have off-target effects. Future studies could address this possibility by identifying I-BET151 bound molecules in trypanosome lysate using mass spectrometry. Future work could also address whether perturbation of individual RNA binding proteins that are altered following I-BET151 treatment in insect stage parasites might recapitulate portions of the differentiation program. More broadly, an increased understanding of gene regulatory mechanisms in this highly diverged eukaryote may shed light on how gene regulatory systems evolved across diverse forms of life.

## Methods

### Parasite Culture and Drug Treatment

Procyclic PF427 parasites were cultured in SDM79 media supplemented with 10% FCS at 27ºC. Parasites were treated with 20µM I-BET151 (Sigma-Aldrich). *EP1/GFP* procyclic reporter parasites were generated as described in^30^. *EP1/GFP* bloodstream parasites^30^ and Single Marker (SM) parasites^77^ (used as controls for verification of the *EP1/GFP* reporter construct) were grown in HMI9 supplemented with 10% FCS and 10% Serum Plus at 37 ºC and 5.0% CO_2_.

### PCR primers

PCR was used to confirm correct integration of the *EP1/GFP* reporter construct using a primer upstream of *EP1* (gtccgataggtatctcttattagtatag) and within GFP (agaagtcgtgctgcttcatgtggt). Primers specific to a region downstream of *EP1* were used as controls (ggccatactagtctttgaatttggatcttaaaattattattg and ggccatctcgagcaacttcagctgcggggc).

### Library Preparation and Sequencing

RNA extractions were performed using RNA Stat-60 (Tel-Test) according to the manufacturer’s instructions. 5µg of RNA was subjected to DNAse treatment and poly(A)^+^ selection using the NEBNext Poly(A) mRNA magnetic isolation module (E7490). High-throughput sequencing libraries were generated using the NEBNext Ultra Directional RNA library prep kit (E7420). Sequencing was performed on an Illumina HiSeq 2000 sequencer using 100bp reads.

### Bioinformatic Analysis

Trimming for quality was performed on fastq files using TrimGalore from Babraham Bioinformatics http://www.bioinformatics.babraham.ac.uk/projects/trim_galore/) with level 3 stringency using the following command (`trim_galore—stringency 3`). Trimmed reads were aligned uniquely to the Tb927v5 reference genome using bowtie allowing 2 mismatches with the following command (`bowtie—best—strata -t -v 2 -a -m 1`). For the VSG analysis reads were aligned to the 427 VSGnome^7^ using the same parameters. Raw counts and RPKM for reads assigned to each gene were calculated using SeqMonk from Babraham Bioinformatics (https://www.bioinformatics.babraham.ac.uk/projects/seqmonk/). The DESeq2^78^ R package was used to identify differentially expressed genes and calculate Benjamini-Hochberg adjusted p-values. The MFuzz R package^32^ was used to cluster differentially expressed genes with a Benjamini-Hochberg adjusted p-value of less than 0.01 using a Likelihood Ratio Test (LRT) analysis. Cluster members with a membership score greater than 0.7 (high core members) are provided in Supplemental Table 6.

Gene Set Enrichment Analysis^79^ was used to identify gene sets that were overrepresented in the set of differentially expressed genes with the following flags:

~~~
-collapse false -mode Max_probe -norm meandiv -nperm 1000 -permute
gene_set -rnd_type no_balance -scoring_scheme weighted -rpt_label
~~~

and

~~~
-sort real -order descending -create_gcts false -create_svgs false
-include_only_symbols true -make_sets true -median false -num 100
-plot_top_x 20 -rnd_seed timestamp -save_rnd_lists false -set_max
500 -set_min 3 -zip_report false
~~~

Gene sets are provided in Supplemental Table 1. Normalized read counts and p-values for differential expression are provided in Supplementary Tables 2 and 3, respectively.

GO analysis on each cluster was performed using TriTrypDB. GO results on Cluster 3 are provided in Supplemental Table 4.

### Flow Cytometry

Flow cytometry on *EP1/GFP* reporter parasites was performed on a Novocyte 2000R from Agilent (formerly Acea biosciences) and analyzed using FloJo software.

