## Supplemental Figures for "Chemical inhibition of bromodomain proteins in insect stage African trypanosomes perturbs silencing of the Variant Surface Glycoprotein repertoire and results in widespread changes in the transcriptome"

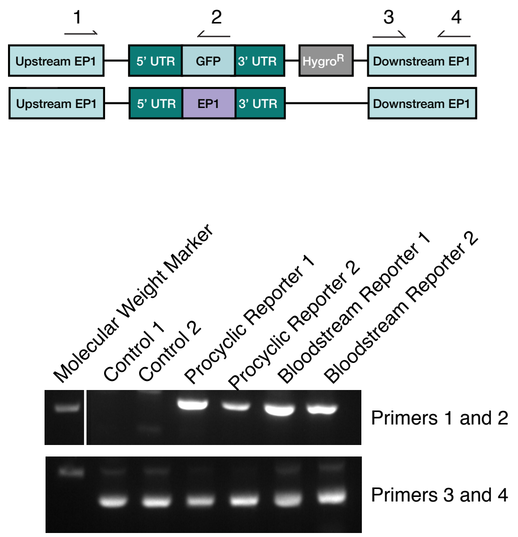


**Supplemental Figure 1.** Top, schematic for *EP1/GFP* reporter construct integrated at the *EP1* locus in procyclic stage parasites. Bottom, PCR confirmation of correct integration of the *EP1/GFP* reporter construct. Genomic DNA was isolated from indicated parasite lines and PCR was performed with primers specific to correct integration (Primers 1 and 2) and control primers (Primers 3 and 4). Control 1 and Control 2 are samples from the Single Marker (SM) bloodstream parasite line that does not contain the *EP1/GFP* reporter construct.

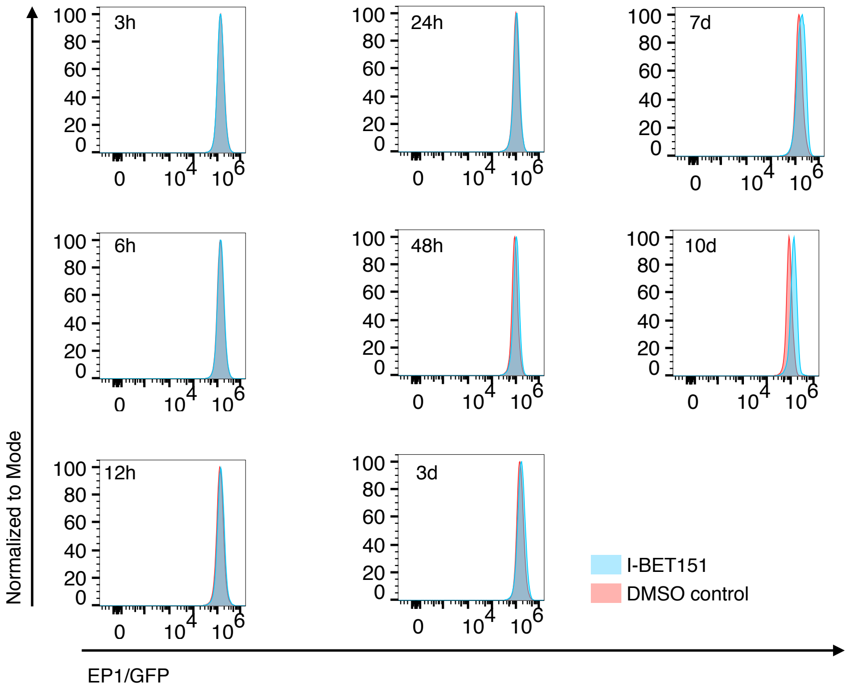


**Supplemental Figure 2**. Flow cytometry for *EP1/GFP* expression in I-BET151 treated parasites for the indicated length of time and compared to a DMSO control.

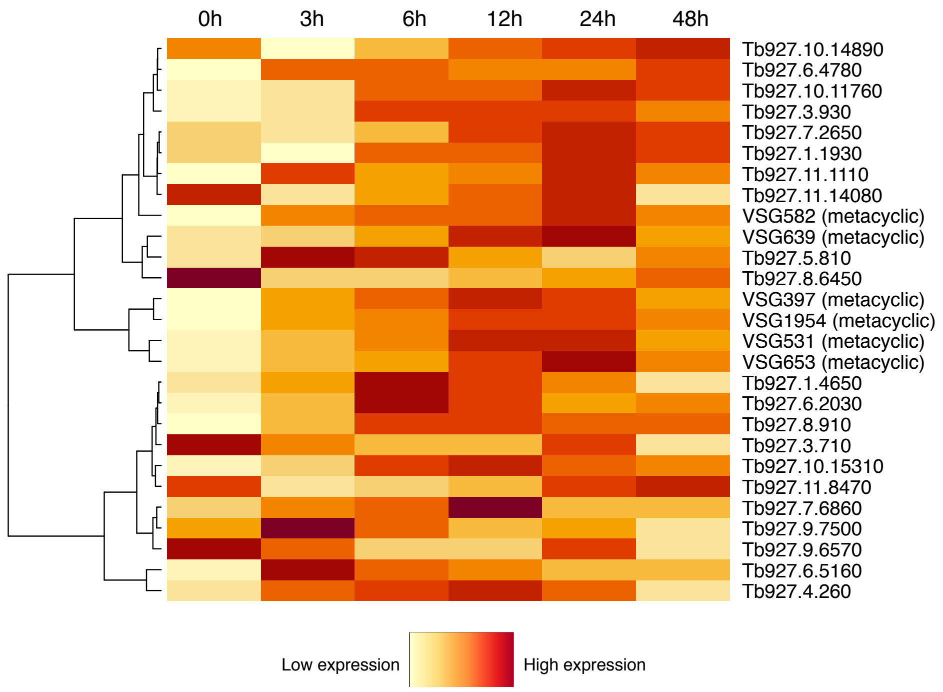


**Supplementary Figure 3.** Heatmap of early metacyclic Meta1 genes as described by Vigneron et al. for procyclic parasite RNA-seq samples at the indicated time of I-BET151 treatment.

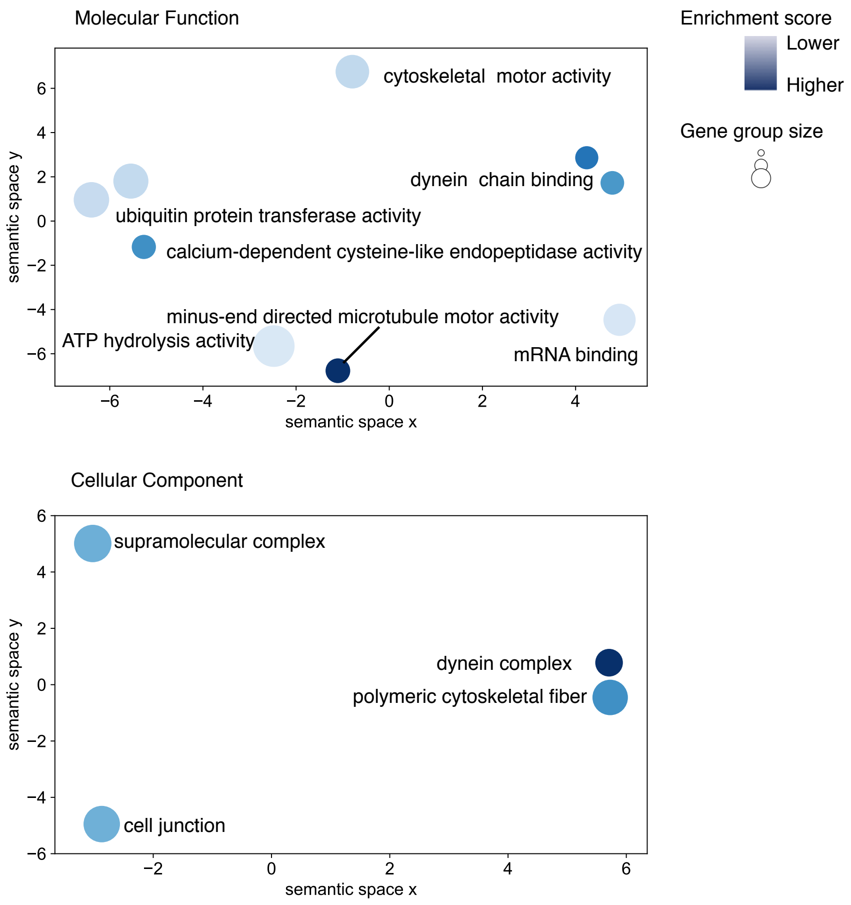


**Supplemental Figure 4.** REVIGO plot of GO enrichment analysis performed on Cluster 3 from the I-BET151 treated dataset.
